# Early-life temperature drives recruitment success in Eurasian perch (*Perca fluviatilis*) populations

**DOI:** 10.1101/2025.11.04.686284

**Authors:** Valentin Cavoy, Jean Guillard, Cécilia Barouillet, Orlane Anneville, Najwa Sharaf, Christian Gillet, Chloé Goulon

## Abstract

Interannual fluctuations in the abundance of young-of-the-year (YOY) European perch (*Perca fluviatilis*) were studied in two large French peri-alpine lakes using a 12-year dataset of late summer hydroacoustic surveys. Previous research based on sampling over only a few years in localized areas or experimental systems has already highlighted the importance of temperature. However, these studies often considered only monthly or seasonal averages. In our study, we sought to understand the factors behind the significant annual fluctuations in perch stock and analysed 12 years of data at the ecosystem level. We used traditional thermal indices based on average values, as well as novel indices reflecting daily temperature fluctuations, which were derived from 36 years of *in situ* perch spawning data. Embryonic thermal conditions alone accounted for between 44% and 88% of the variation in YOY density across the lakes. An increase in temperature during the embryonic phase was beneficial, whereas significant daily variations had the opposite effect. In contrast, trophic factors and the adult stock did not exhibit consistent trends. Our study confirms previous findings using ecosystem-scale analyses, long-term data, and novel thermal indices, validating *in natura* the central role of spring thermal regimes in shaping recruitment and improving our understanding of recruitment variability under climate change.

## 1 INTRODUCTION

Interannual variability in fish recruitment remains one of the most enduring challenges in ecology (Houde, 2008; Takasuka et al., 2021). In both marine and freshwater systems, explaining and predicting year-class strength is difficult because of the complex interplay between environmental and biological drivers (Brucet et al., 2013; Houde, 2008; Marcek et al., 2021). Despite decades of research, several influential conceptual frameworks, including the critical period (Hjort, 1914), match–mismatch (Cushing, 1990) and larval size hypotheses (Miller et al., 1988), have not yielded consistent predictive power across systems. However, a common thread uniting these hypotheses is their fundamental reliance on temperature-driven processes. Temperature governs nearly every stage of early development in fishes. It controls the rate of embryogenesis, hatching timing, yolk absorption, larval growth and metabolism (Ayala et al., 2015; Ojanguren & Braña, 2003). It also determines the duration and onset of the transition from endogenous to exogenous feeding, a critical bottleneck in recruitment, as in clupeid species such as herring (Houde, 2008).

Temperature can also indirectly affect fish recruitment. The match–mismatch hypothesis, for instance, is ultimately temperature-dependent: larval hatching must align with the phenology of zooplankton growth, which is governed by thermal cues (Cushing, 1990; Peck et al., 2012). Similarly, the critical period hypothesis hinges on the pace of developmental progression and the metabolic demands imposed by temperature (Ayala et al., 2015; Ojanguren & Braña, 2003). Together, these frameworks converge on the idea that temperature is not merely one variable among others but is the structuring force that shapes the timing, success and synchrony of early-life-stage processes (Laurel et al., 2021).

For Eurasian perch (*Perca fluviatilis*), early development occurs during a short and thermally sensitive spring window (Craig, 1987; Gillet & Dubois, 1995, 2007). The direct effect of temperature during early life stages on the recruitment of both *P. fluviatilis* and *P. flavescens*, two species that are morphologically and ecologically similar (Thorpe, 2011), have been investigated since the 1980s (Fraz et al., 2024, 2025; Guma’A, 1978; Linløkken, 2023; Ning et al., 2025; Schmitz & Sepulveda Villet, 2021; Wang & Eckmann, 1994). Early developmental stages thus remain highly sensitive to thermal conditions. For example, a wind-induced decrease in surface water temperatures in spring can delay warming trends and cause high mortality (Clady, 1976; Van De Hey et al., 2013).

However, temperature is not the only factor affecting early perch development (Craig, 2000). Wind can act as a mechanical stressor by physically disrupting the egg ribbons laid in shallow areas, displacing or damaging them and reducing hatching success (Clady & Hutchinson, 1975). In parallel, biotic factors such as zooplankton density, spawner abundance, and cannibalism add further complexity to recruitment dynamics (Houde, 2008). Zooplankton availability is particularly critical for larval perch, which are selective feeders that require appropriately sized prey during a narrow posthatching window (Wang, 1994). Moreover, adult perch density can have both positive effects through increased reproductive output and negative effects via intraspecific predation and competition among perch juveniles (Craig, 2000; Dubois *et al*., 2008; Kokkonen et al., 2019).

Previous studies have mostly assessed the effects of temperature under controlled conditions, typically in experimental systems at constant temperatures (Fraz et al., 2024, 2025; Guma’A, 1978; Van De Hey et al., 2013). Although other studies have been conducted on natural ecosystems, they have relied on point-in-time indicators of perch density, such as bottom trawl surveys, gillnets and traps, which are not representative of the ecosystem on a large scale, or on commercial fishing landings, which are known to be biased (Clady, 1976; Clady & Hutchinson, 1975; Honsey et al., 2016). Furthermore, these studies often compare young-of-the-year (YOY) density or aquarium survival to broad-spectrum indicators such as the mean spring–summer temperature (Clady, 1976; Honsey et al., 2016). We have noted a lack of studies aimed at validating these results over long periods of time in natural ecosystems where the entire environment was monitored. Similarly, we found very few studies that used high-resolution thermal indices capable of capturing short-term temperature fluctuations during early developmental stages, despite growing evidence that temperature variability can strongly influence biological responses (Vasseur et al., 2014).

In this study, we assessed the influence of key environmental and biological drivers, focusing primarily on temperature as a stage-specific, phenology-aligned predictor of recruitment in Eurasian perch. Daily temperature fluctuations during incubation and larval development result in a complex thermal profile, which is difficult to reduce to a single representative value. To resolve this, we tested several thermal indices, each capturing different aspects of the thermal regime, to identify the one that best explains the fluctuations in YOY perch density in lakes. We hypothesized that compared with conventional seasonal temperature means, dynamic temperature metrics, which are based on the biological timing of early development, are stronger predictors of YOY perch abundance. To test this hypothesis, we used YOY density proxies derived from hydroacoustic surveys conducted between 2011 and 2023, which represent ten effective years of data, combined with environmental and biological monitoring in two deep French peri-alpine lakes, Annecy and Bourget (Rimet et al., 2020). Recruitment variability was then related to indices describing water temperature, wind speed, zooplankton availability, and adult perch abundance.

## 2 MATERIALS AND METHODS

### 2.1 Study sites

Lake Annecy (45°53’N, 6°08’E) and Lake Bourget (45°43’N, 5°52’E) are two geographically close French peri-alpine lakes (Table 1). They possess similar physical characteristics (e.g., size and maximum and average depths) but differ in terms of trophic dynamics, biodiversity and fisheries management (Changeux et al., 2024; Jacquet et al., 2014). Lake Annecy is oligotrophic, with total phosphorus concentrations being less than 10 µg P.L^-1^ since 1994 (Frossard et al., 2024). Lake Bourget experienced a period of strong eutrophication (120 µg P L⁻¹ in 1983) followed by a reoligotrophication phase, and since the early 2020s, most indicators classify it as oligotrophic (Jenny et al., 2024). The phosphorus concentrations from recent years are shown in Supplementary Figure 1.

**Table 1:**
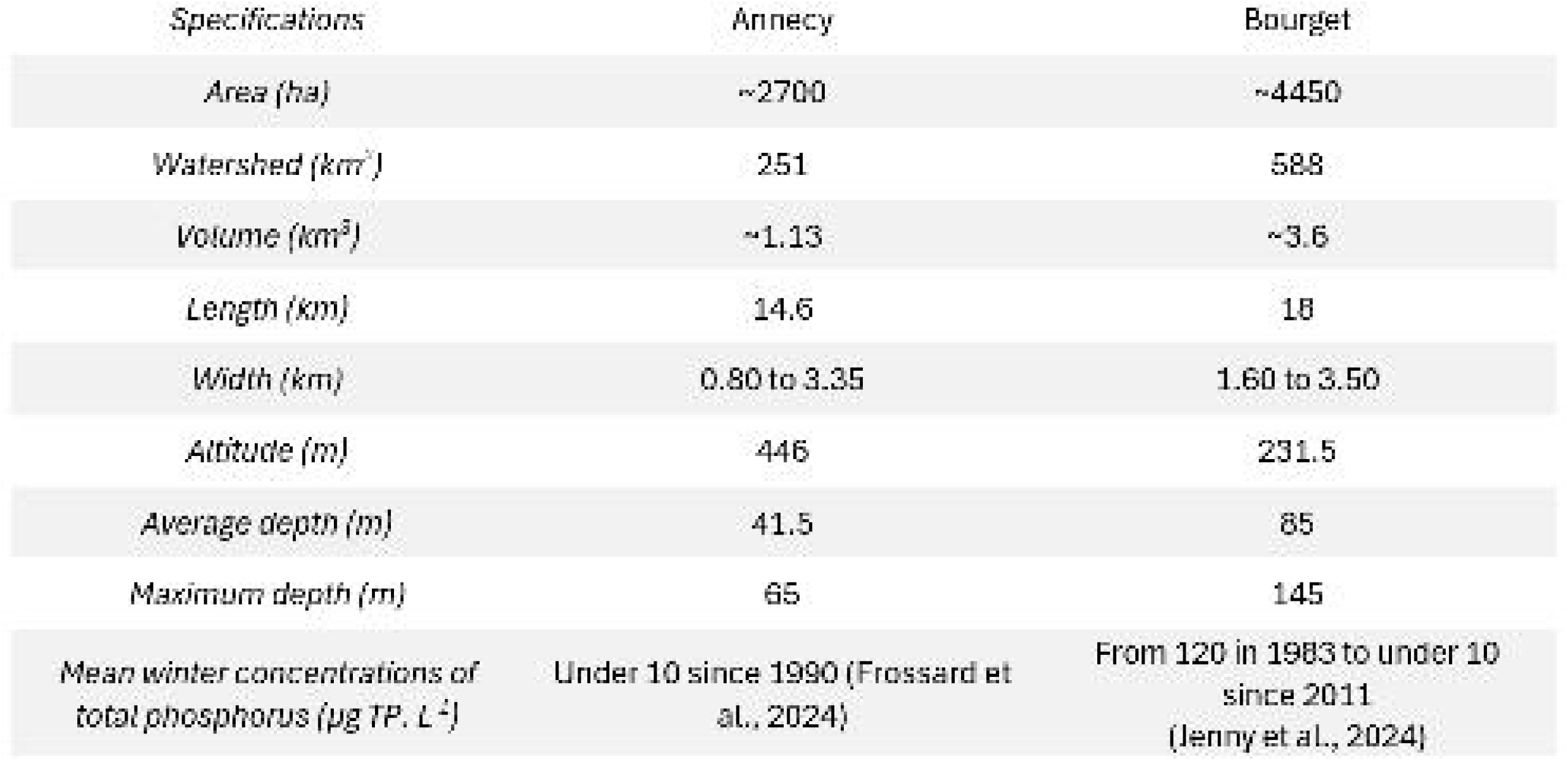
Characteristics of the study lakes.

Approximately fifteen fish species are found in Lake Annecy, twenty in Lake Bourget, and perch is the numerically dominant species in both lakes (Frossard et al., 2024; Jenny et al., 2024). Based on standardized gillnetting surveys (OLA-IS, AnaEE-France, INRAE of Thonon-les-Bains, SILA, CISALB, (Rimet et al., 2020), European perch accounted for, on average (and across all age classes), 83% of the individuals captured in both the benthic and pelagic gillnets in Lake Annecy and 66% in Lake Bourget (Frossard et al., 2024; Jenny et al., 2024).

### 2.2 Estimation of YOY perch density

Following standard protocols (CEN, 2014) and recommendations (Draštík et al., 2017), hydroacoustic surveys were performed annually at night during late summer (mid-September for Lake Annecy and end of September to early October for Lake Bourget) from 2011– to 2023 using scientific echo-sounders (Guillard et al., 2006). Initially, a split-beam EK60 Simrad (70 kHz; Simrad Kongsberg Maritime AS, Horten, Norway) was used, and most recently, surveys were performed with a split-beam EK80 Simrad (120 kHz). Hydroacoustic data collected using these two frequencies and generations of sounders yield comparable results (De Robertis et al., 2019; Guillard et al., 2014; Mouget et al., 2019; Rautureau et al., 2022). Detailed protocols are provided in Guillard et al. (2006) for Lake Annecy and in Yule et al. (2013) for Lake Bourget, as well as in the annual survey reports for both lakes (Frossard et al., 2024; Jenny et al., 2024). These surveys provide an indicator of fish density (number of fish. ha^-1^) in the upper water layer above the thermocline, where YOY perch predominantly resides (Guillard et al., 2006; Yule et al., 2013) but also, to a lesser extent, other species such as young roach. It is not possible to identify the species of fish detected by hydroacoustics; however, in scientific pelagic gillnet surveys conducted during the same period, YOY perch consistently accounted for most fish caught above the thermocline, averaging 93% in Lake Annecy and 74% in Lake Bourget (OLA-IS, AnaEE-France, INRAE Thonon-les-Bains, SILA, CISALB, Rimet et al., 2020, Frossard et al., 2024; Jenny et al., 2024). Although these proportions fluctuated in years with low overall pelagic catches, the median values remained high at 96% in Annecy and 77% in Bourget. Consequently, fish density estimates from these surveys provide a reliable proxy for the annual strength of the YOY perch cohort throughout the studied period (Guillard et al., 2006; Sotton et al., 2011), referred to in the text as YOY perch density.

### 2.3 Determination of developmental stages

To investigate the relationships between environmental variables and YOY perch density, we defined the duration of the critical stages of the perch life cycle and selected the main environmental variables influencing perch density during this period based on the literature (Craig, 2000; Fraz et al., 2024; Gillet & Dubois, 2007; Schmitz & Sepulveda Villet, 2021; Wang & Eckmann, 1994). The timing of the early developmental stages of perch in Lakes Annecy and Bourget was estimated from the spawning period in Lake Geneva. These three lakes are geographically close, with Lake Geneva located approximately 40 km from Lake Annecy and 60 km from Lake Bourget. The timing of early development was also inferred from degree days (°C Day), defined as the cumulative sum of daily mean temperatures above 0 °C, which is used to quantify developmental rates in fishes (Neuheimer & Taggart, 2007).

In peri-alpine lakes, perch spawning is triggered by water temperature in the littoral zone (Gillet & Dubois, 2007). For instance, in Lake Geneva, the date at which 50% of eggs are deposited is correlated with the date at which the surface water temperature reaches 12 °C (Gillet & Dubois, 2007). Based on the method described in Gillet and Dubois (2007), the phenology of perch reproduction in Lake Geneva has been studied continuously since 1986 (Goulon et al., 2024).

For each year, records of spawning dates and the associated number of eggs laid are available for Lake Geneva, along with daily modelled surface water temperatures (Sharaf et al., 2023). In this study, we analysed 36 years of perch monitoring records and found that egg laying is concentrated between 10.5 °C and 13.3 °C, which are the temperatures at which 10% and 90% of the total spawning events, respectively, have occurred (Supplementary Figure 2). The maximum spawning activity occurred at 11.5 °C. Based on this analysis, we estimated the spawning periods of perch in peri-alpine lakes as the dates when the surface water temperature reached 10.5 °C (10% of the number of eggs deposited) and 13.3 °C (90% of the number of eggs deposited). These temperature thresholds are valid only for lakes within the same geographical region, as perch spawning temperatures vary from north to south across the geographical distribution of the species (Thorpe, 2011).

Considering incubation periods of 13−14 days at 14 °C and 15−16 days at 12 °C (Guma’A, 1978), perch eggs would require 187.5 ± 8 degree days. The threshold was set at 187.5 degree-days, and the duration of the larval stage was defined as 33 days after hatching based on literature values (Vlavonou, 1996).

### 2.4 Parameters considered for the embryonic stage

The embryonic stage begins when the surface temperature reaches 10.5 °C, which corresponds to the start of the spawning period (Figure 1). Even if the first eggs laid become larvae approximately 187.5 degree-days after the date of spawning, the embryonic phase does not stop at that point. To consider the theoretical incubation phase of all the spawns, we identified the time at which the incubation phase of the last eggs laid ended, i.e., 187.5 °C days after a temperature of 13.3 °C was reached for surface water. The different dates for the beginning and end of the embryonic and larval periods for each lake and each year are summarized in Supplementary Table 1.

**Figure 1:**
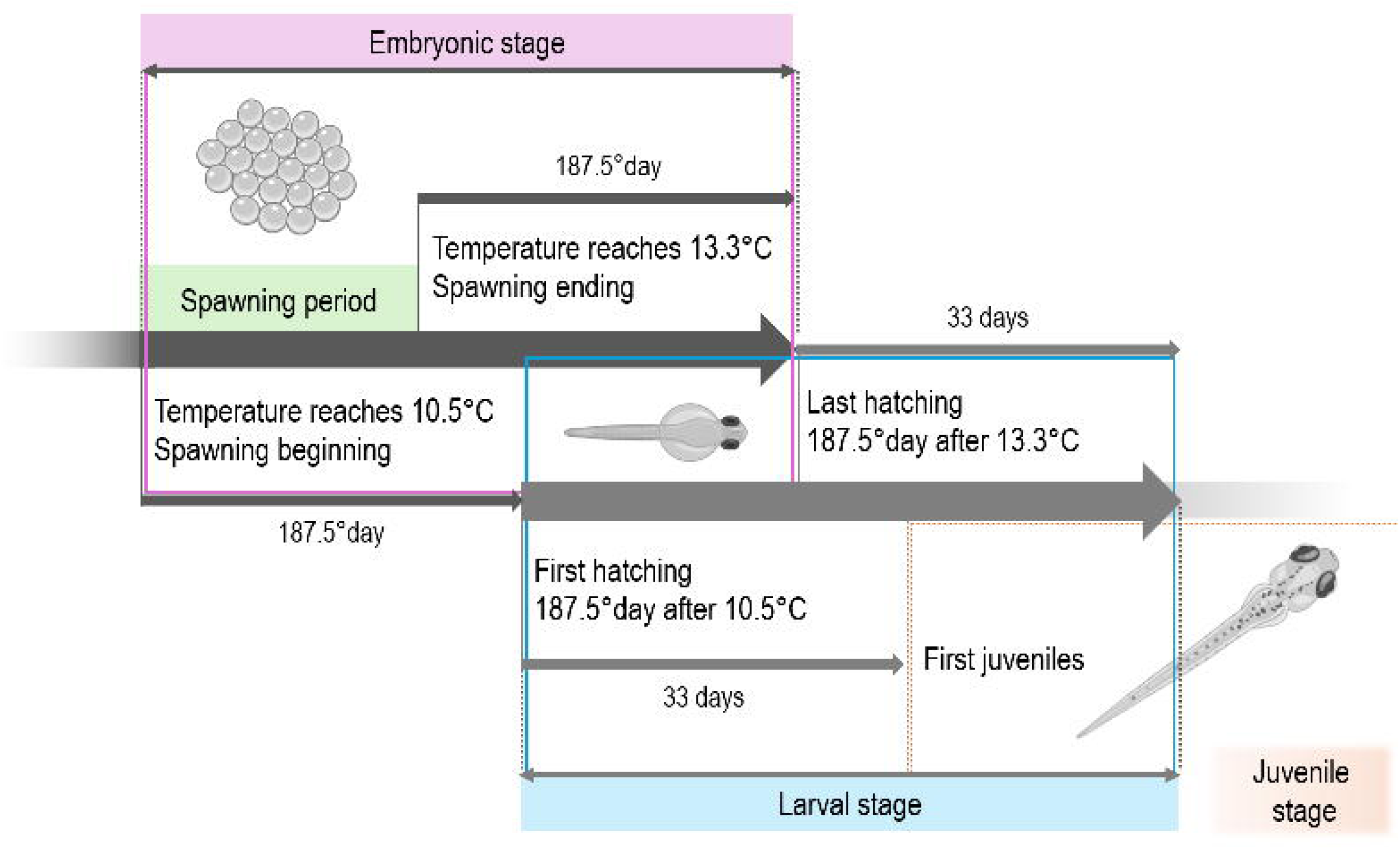
Summary diagram of the early developmental stages of perch based on our survey data and the literature (Guma’A, 1978; Vlavonou, 1996).

**Figure 2:**
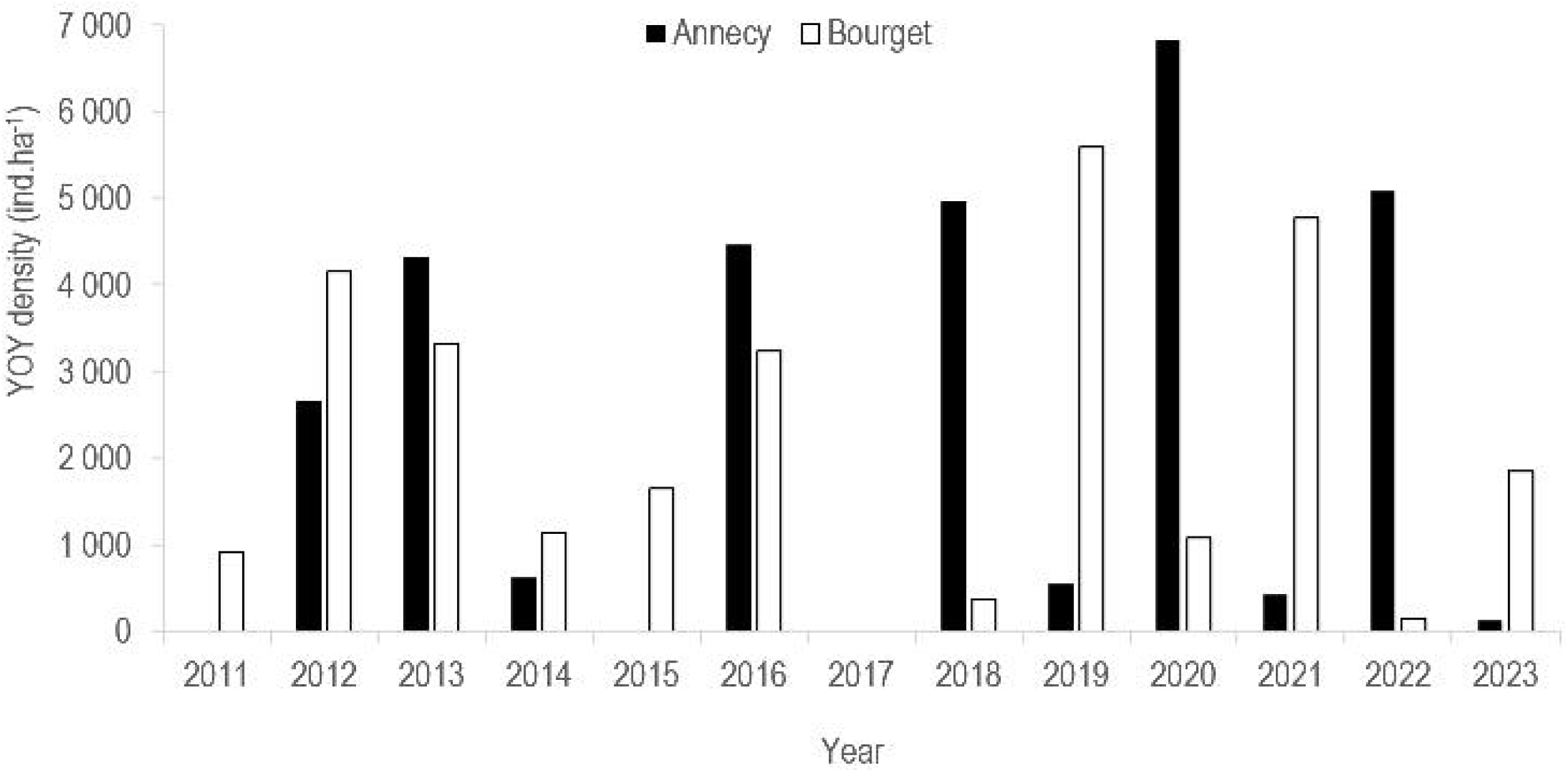
Annual YOY perch density (number of fish detected per hectare, estimated by hydroacoustics) in Lakes Annecy and Bourget.

#### 2.4.a Temperature data and thermal indices

*In situ* surface temperature data were acquired using Tinytag or Hobo Onset underwater data loggers between 2011 and 2023 for Lakes Annecy and Bourget, respectively. In Lake Annecy, temperatures were measured in the littoral zone every half-hour at 4 stations between 0.5 and 1 m depth (data from the SILA, Syndicat Mixte du Lac d’Annecy – Lake Annecy Mixed Syndicate). In Lake Bourget, measurements were recorded every minute at a single monitoring station at a depth of 1.2 m with a continuous temperature record covering the entire study period (data from the CISALB, Comité Intercommunautaire pour l’Assainissement du Lac du Bourget – Intercommunal Committee for the Sanitation of Lake Bourget). Although additional stations were available in Lake Bourget, the time series were not long enough and were not included. Nevertheless, surface temperatures were comparable among the stations within each lake, with no significant differences observed. Furthermore, since perch generally reproduce in coastal habitats, most often in the top few meters of the water column (Gillet & Dubois, 2007), the temperature measurements used here are highly representative of spawning conditions.

The mean daily surface water temperature was calculated at each station and subsequently averaged across the stations for Lake Annecy. In Annecy, the number of active monitoring stations has varied over time, with between one and four measurements available per day depending on the operational conditions. For periods with missing *in situ* measurements (e.g., during probe maintenance, vandalism, and data loss), which represented 25% of the April–June sequences in Annecy and 7% in Bourget (Supplementary Figure 3), daily surface temperatures were reconstructed using simulations from the two-layer semiempirical Ottosson–Kettle–Prats Lake Model (OKPLM) (Prats & Danis, 2019; Sharaf et al., 2023). The model simulates temperatures based on the geomorphological characteristics of the lake as input and is forced with meteorological data, including air temperature and solar radiation. In this study, simulations were conducted using model parameters calibrated with observation data from the Observatory of LAkes (OLA) (Rimet et al., 2020) and meteorological data from SAFRAN, Système d’Analyse Fournissant des Renseignements Adaptés à la Nivologie – Analysis System Providing Information Tailored to Snow Science (Durand et al., 1993).

Based on these surface temperature data, we calculated four indices to characterize the thermal conditions relevant to embryonic development. The first index, the mean embryonic thermal index (MET), corresponds to the mean temperature of the embryonic phase and captures the overall thermal conditions known to influence egg development rates in perch (Guma’A, 1978; Hokanson & Kleiner, 1974; Linløkken, 2023). The second index, the embryonic thermal amplitude index (ETA), represents the thermal amplitude during incubation and is calculated as the difference between the highest and lowest daily mean temperatures, reflecting the potential impact of thermal variability on embryonic survival (Schmitz & Sepulveda Villet, 2021; Van De Hey et al., 2013).

The third index, the embryonic thermal index (ETI), was specifically developed in this study to account for differences in daily mean temperature during egg incubation, allowing discrimination between unfavourable years (i.e., stagnant and/or highly variable temperatures) and optimal years (i.e., regular warming of water). This index is based on the known sensitivity of fish embryos to short-term temperature fluctuations (Clady, 1976; Clady & Hutchinson, 1975; Schmitz & Sepulveda Villet, 2021; Vasseur et al., 2014). The ETI is calculated as follows:

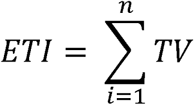

where temperature variation (TV) corresponds to the difference between two consecutive daily mean temperatures in °C, with one value each day from the first to the last day of incubation (n). The reliability of the ETI calculation based on fixed temperature thresholds was assessed using the observed spawning phenology from Lake Geneva (see Supplementary Sections 5 and 6).

Finally, the thermal embryonic variation index (TEV) is derived from the ETI using absolute temperature variation values and is calculated as follows:

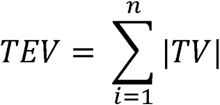

where |*TV*| is the absolute value of temperature variation in °C, with one value each day from the first to the last day of incubation (n), providing a measure of cumulative thermal instability during incubation.

#### 2.4.b Wind speed index

During the embryonic phase, in addition to temperature, wind is considered a potential factor influencing incubation success (Bourinet et al., 2023; Clady & Hutchinson, 1975). To capture its influence, the embryonic wind condition index (EWC) was defined as the mean wind speed (m/s) during the embryonic phase. For additional information on the wind speed data, refer to the Supporting Information (Section S7).

### 2.5 Parameters considered for the larval phase

The larval phase partially overlaps with the embryonic phase; as the first larva emerges from the eggs, other embryos are still developing (Figure 1). The larval phase thus begins 187.5 days after the surface temperature reaches 10.5 °C, corresponding to the first hatch, and ends 33 days after the surface temperature reaches the threshold of 13.3 °C plus 187.5 °C for hatching, corresponding to the date when the last larva reaches the juvenile stage. The data are summarized in Supplementary Table 1.

#### 2.5.a Temperature data and indices

After hatching, larvae remain strongly influenced by temperature, which impacts individual growth (Craig, 2000; Linløkken, 2023). The four thermal indices originally used for the embryonic phase were adapted for the larval phase. The mean larval thermal index (MLT) corresponds to the mean temperature during the larval phase, and the larval thermal amplitude index (LTA) represents the difference between the maximum and minimum temperatures during larval development. Like the ETI, the larval thermal index (LTI) sums the daily temperature variation (TV) during the larval phase:

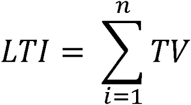

Finally, the thermal larval variation index (TLV) is similar to the LTI but uses absolute values of TV to represent the total thermal variations experienced by the larvae during their 33 days of larval development:

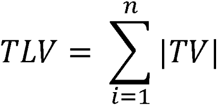

#### 2.5.b Other environmental parameters

In addition to temperature, other environmental parameters were considered for the larval phase. These include the larval wind condition index (LWC), mean wind speed during the larval period, zooplankton prey availability at first feeding indices (i.e., rotifers, zai_rot; copepods zai_cop; Cladocera, zai_clad; and all available prey zai_all) (Craig, 2000; Masson et al., 2001; Wang, 1994) and adult perch abundance as genitor stock indicator (Gen index) and as intraspecific competition or predation proxies (Pred index) (Craig, 1978, 2000; Gillet & Dubois, 2009). Detailed descriptions of these parameters are provided in the Supporting Information (S8 and S9 section).

To assess the lake trophic dynamics and nutrient availability, the mean concentrations of total phosphorus (TP) and orthophosphate in the 0–15 m water layer from April to June (2011–2023) were also examined. The TP and orthophosphate concentrations were obtained from OLA survey data (Rimet et al., 2020) and correspond to depth-weighted means expressed as mg P L⁻¹ (elemental phosphorus), determined by the molybdenum-blue method after sulfuric acid digestion (EPA 365.3). For clarity, dissolved inorganic phosphorus concentrations from orthophosphate are reported as µg P L⁻¹ throughout this study. These nutrient concentrations can serve as proxies for nutrient availability (Schindler, 1978), which is expected to influence the spring development of the primary trophic network, support zooplankton production, and ultimately sustain *Perca fluviatilis* larvae during their critical early life stages (Wang & Eckmann, 1994). During this period, the TP concentrations fluctuated between 4 and 9 µg L^-1^ in Lake Annecy and between 7 and 17 µg L^-1^ in Lake Bourget (Supplementary Figure 1).

### 2.6 Statistical analysis

Statistical analyses were performed in RStudio, Version 2025.05.0, using the R statistical environment (R Core Team, 2025). Generalized additive models (GAMs) were used to model the nonlinear functional relationship between our set of explanatory variables and the annual YOY perch density for each lake using the mgcv (Wood, 2022) and gratia (Simpson & Singmann, 2022) packages, following the methods described in Simpson and Anderson (2009). Given the number of observations available, to avoid overlearning and not excessively decrease our degrees of freedom, simple covariate models were developed. Each GAM was modelled using thin-plate regression splines as the basis spline, and the degree of smoothness was determined via restricted maximum likelihood (REML). A total of eighteen explanatory variables were tested (Table 2). The model’s residuals were checked using the *appraise* function from the *gratia* package (Simpson, 2024), the *pacf* function and the associated mathematical tests (Shapiro, Breusch□Pagan, Durbin-Watson). For the environmental factors that were identified as significant in both studied lakes (*p* value of the GAM < 0.05), hierarchical GAMs (HGAMs) were subsequently used to assess whether the YOY perch responded similarly to these factors in both lakes (Pedersen et al., 2019). The two datasets were combined, resulting in a total of 22 observations, thus highlighting a possible similar effect of the environmental factors on the YOY perch population abundances of both lakes. To this end, different types of models (G, S, GS, I and GI) were compared to determine the best fit using the method described in Pedersen et al. (2019). The equations used for each model are shown in Table 3. Two categories of models were applied: nonglobal trend models (S and I models) and shared global trend models (G, GS and GI models). For the nonglobal trend models, the “S” model assumes that the lake-specific smoothers have the same wiggleness, whereas for the “I” model, both trends and wiggleness are lake specific. Note that for the latest model, the tensor product allows each of the smoothers to have a lake-specific wiggleness because the “Lake” variable appears to have a random effect structure (bs = “re”). Similarly, for the models with a shared global trend, the “G” model includes only one global smoother for both lakes, whereas the “GS model” includes a common smoother as well as a lake-specific smoother that has the same wiggleness. In the formula of the GS model, the explicative variable (bs = “tp”) corresponds to the common smoother as for the G model, and s(explicative variable, Lake, bs = c (”tp”,”fs”)) corresponds to the lake-specific smoothers. Finally, the “GI model” combines the global trend smoother with a group-level smoother where lakes can have specific smoothers with different wiggleness values. For this most complex model, the formula is s(explicative variable, bs = “tp”), which is a primary parameter for the shared trend; another s(explicative variable, by = Lake, bs = c (”tp”, “fs”)) corresponds to a second lake-scale smoother and a final one s(Lake, bs = “re”), which allows the wiggleness to be adapted to each lake by introducing the Lake variable as having a random effect structure.

**Table 2:**
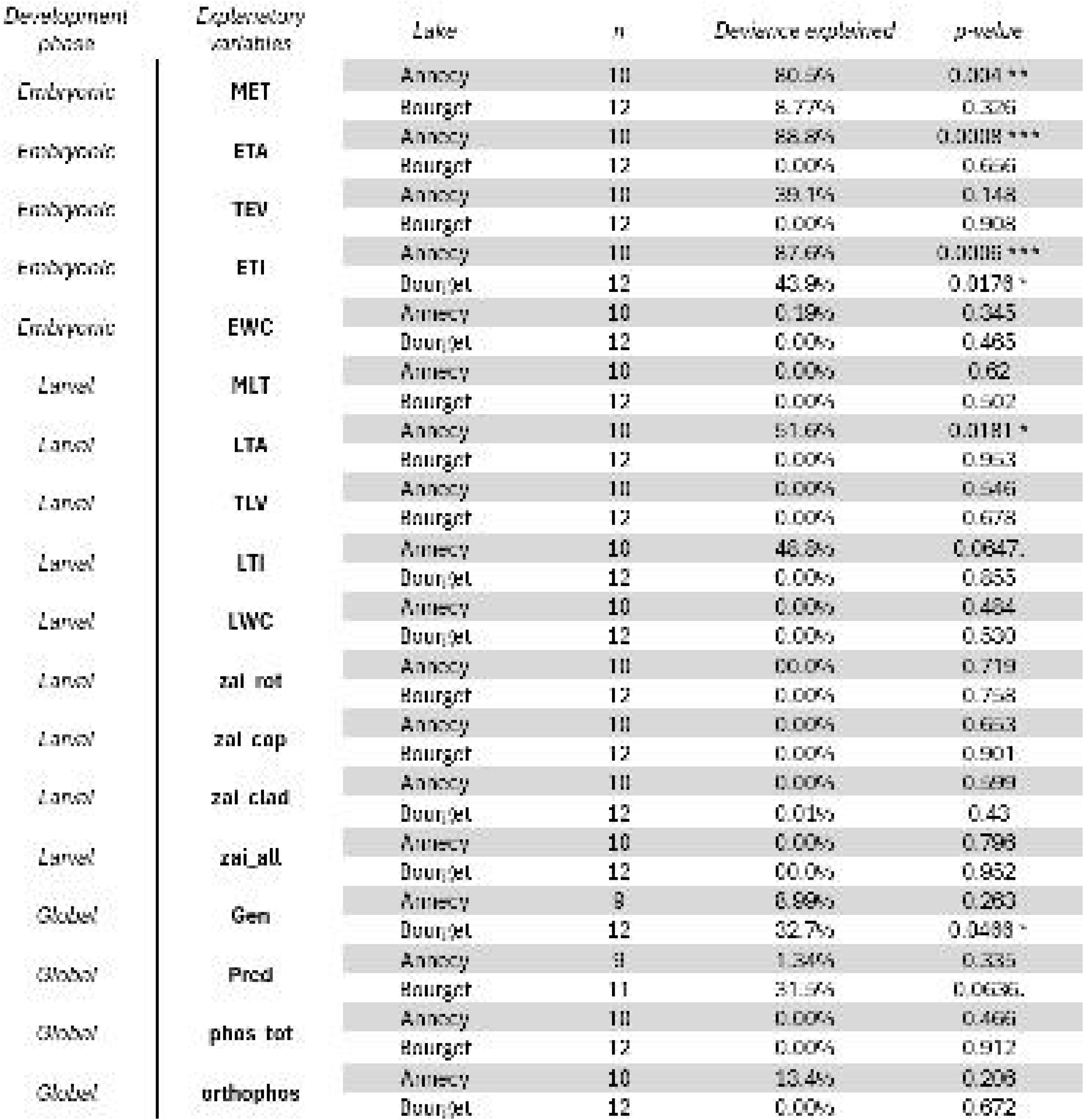
Single covariate GAMs results for the eighteen variables: mean thermal index (MET), embryonic thermal amplitude (ETA), embryonic thermal index (ETI), thermal embryonic variation (TEV), embryonic wind condition (EWC), mean larval thermal index (MLT), larval thermal amplitude (LTA), larval thermal index (LTI), thermal larval variation (TLV), larval wind condition (LWC), zooplankton abundance indeces (rotifers, zai_rot; copepods, zai_cop; cladocerans, zai_clad; and all prey, zai_all), genitor stock index (Gen), predation index (Pred), total phosphorus index (phos_tot) and orthophosphate index (orthophos). For each model, the lake dataset used was specified, as were the number of statistical individuals (n), the explained variance, the p value of the model (signif. codes: *** p < 0.001; ** p < 0.01; * p < 0.05.; *p* < 0.1) and the development phase related to the index.

**Table 3:**
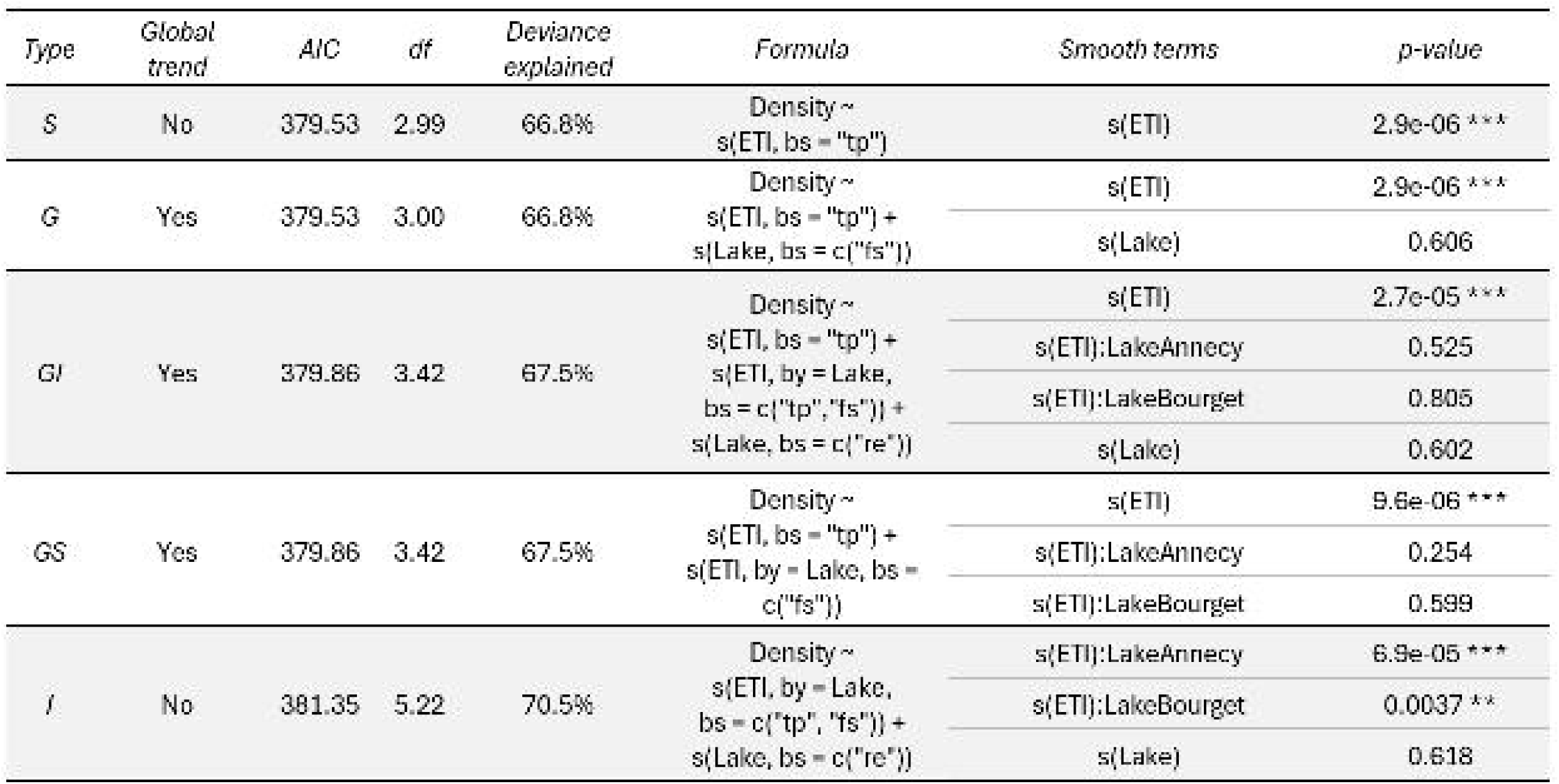
HGAMs results with the combined data from the two lakes. For each model (Pedersen et al., 2019), the global trend, AIC, degree of freedom (df), deviance explained, formula and *p* value by smooth terms (signif. codes: *** *p* < 0.001; ** *p* < 0.01; * *p* < 0.05.; *p* < 0.1) are detailed.

### 2.7 Ethical statement

The utilization of experimental animals was deemed superfluous in the present study. The data collected were obtained from measurements taken in the natural environment on natural fish populations not affected by the study. The use of hydroacoustics, a nonintrusive method, made this possible (more details are provided in Section 2.2).

## 3 RESULTS

### 3.1 Annual YOY perch density

YOY perch densities can vary greatly from year to year and from lake to lake. The minima were recorded at 128 individuals per hectare in Annecy in 2023 and at 149 in Bourget in 2022. The maxima were 5,585 individuals per hectare for Annecy in 2019 and 6,833 for Bourget in 2020. On average, there was a tenfold difference between high and low YOY density years.

### 3.2 Indices by lake

Simple GAMs using a single covariate were processed with the formula: Density (of YOY perch) ∼ s(Explicative variable, bs = “tp”). The results of the GAMs are shown in Table 2. Among the 18 factors per lake model combinations tested, only six were found to have a significant effect on YOY perch density, and five were related to temperature.

In Lake Annecy, 4 factors influenced the YOY density (Table 2), and 3 were significant and related to temperature during the embryonic phase, namely, the MET, the ETA and the ETI. With respect to these three factors, the GAMs explained 80.5%, 88.8% and 87.6% of the deviance, respectively. All three had a similar trend. The higher the explanatory variable is, the higher the density of YOY perch in late summer (Figure 3). However, for ETA, the conditions for homoscedasticity of the model’s residuals were not satisfied; as such, the model of YOY density using the ETA factor could not be validated. During the larval development stage, another significant thermal index, LTA, was found to significantly influence YOY density variability during the larval phase. LTA explained 51.6% of the variability in density. The higher the LTA is, the lower the density.

**Figure 3:**
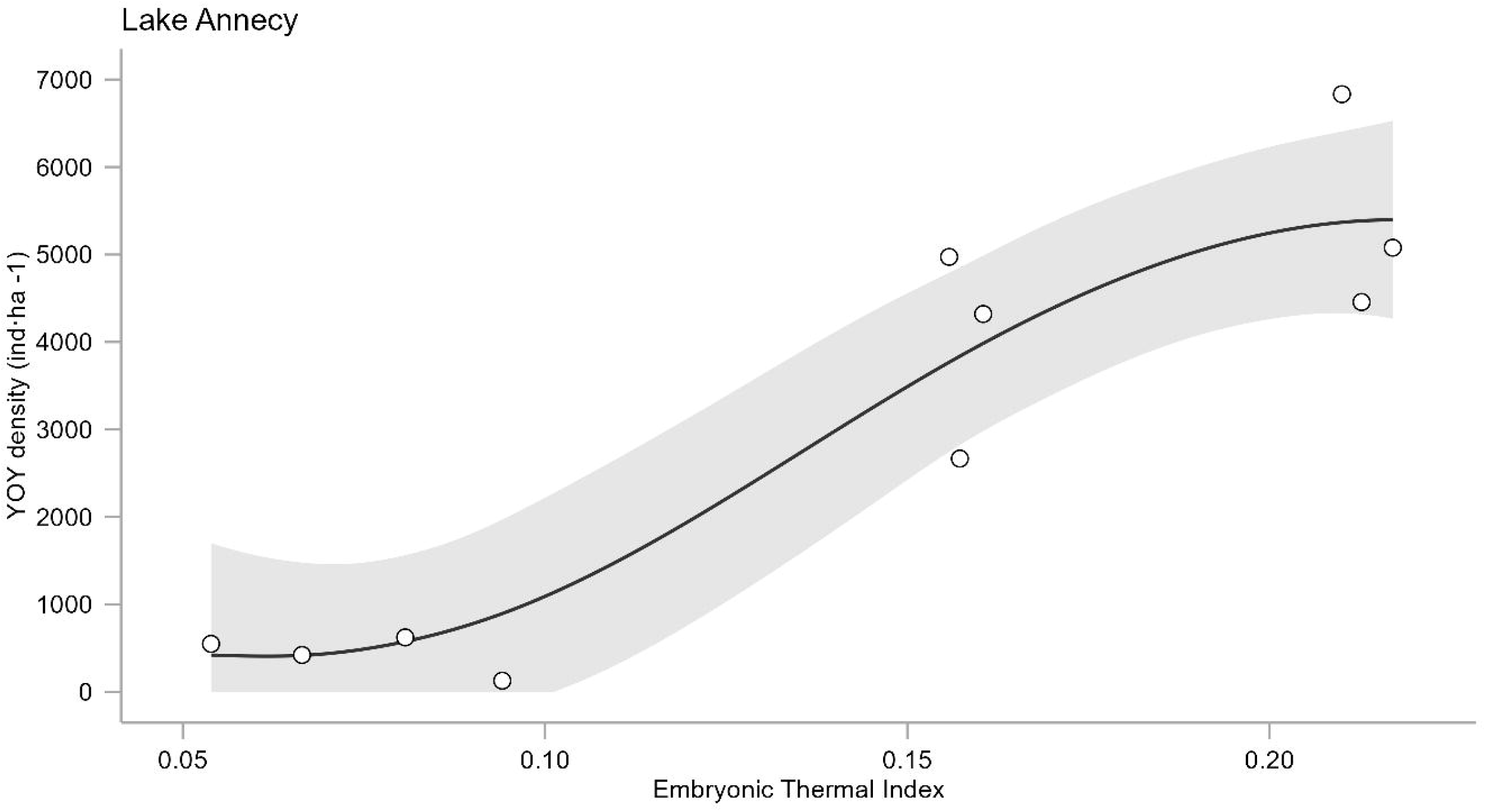
Effects of the embryonic thermal index (ETI) on YOY density in Lake Annecy (Top) and Lake Bourget (Down), single covariate GAMs.

Among the three models found to be relevant for Lake Bourget, one involved a temperature-based predictor during the embryonic stage (Table 2). Indeed, the ETI explained 43.9% of the variation in density for Lake Bourget. The variance explained was lower than that explained in Lake Annecy, but the relationships were similar. The higher the ETI is, the more YOY perch are present at the end of the summer (Figure 3). Another factor not related to the temperature index, the Gen index of the genitor stock, was also significant in Bourget. Explaining 32.7% of the variation in density, the relationship seemed to indicate that when more genitors were present, fewer YOY perch were found at the end of the summer (Supplementary Figure 5). Conversely, the intraspecific predation index, Pred, was not significant (*p* value = 0.0636) but shows the opposite relationship compared with the Gen index. Indeed, the higher the potential cannibalism, the higher the density of YOY perch.

### 3.3 Comparison of the embryonic thermal index tendencies of the two lakes

The only index that was found to be significant and similar for both lakes separately was the ETI. Therefore, the next step was to assess whether the relationship between incubation temperature and YOY perch density at the end of the summer was identical in both lakes or varied depending on the lake. The close performances of these models, particularly G (Figure 4), GS and GI, which all incorporate a global trend component, suggest that a general positive relationship between embryonic thermal conditions (ETI) and YOY perch density is shared between the two systems. This interpretation is supported by the overall pattern observed across the models: YOY density consistently increased with increasing ETI values in both lakes. The model GI, which includes lake-specific smoothers, and model I, which fits entirely separate responses for each lake, were slightly less parsimonious, indicating that although local particularities may exist, the dominant effect of embryonic temperature does not significantly differ between lakes.

**Figure 4:**
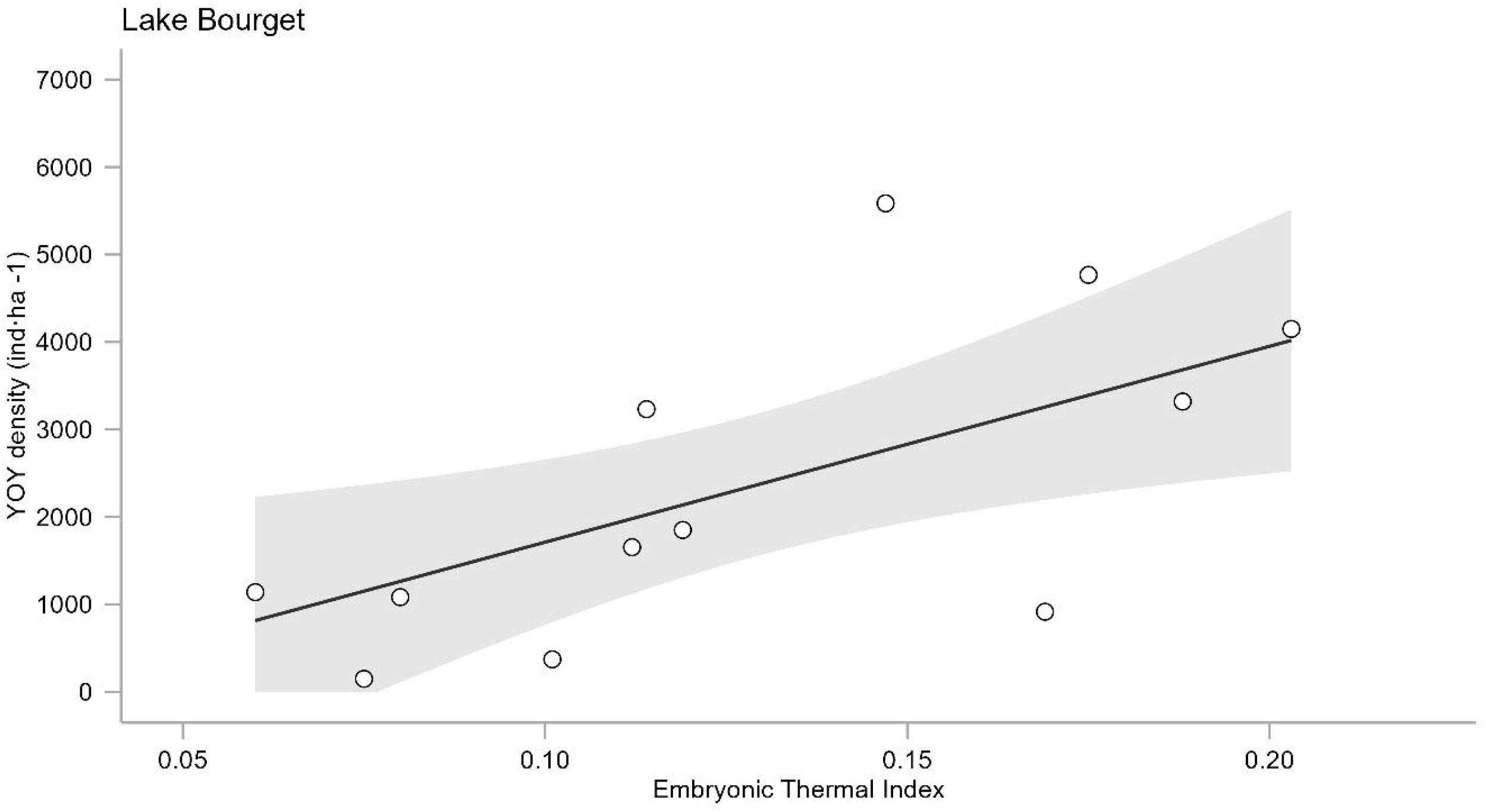
Effect of the embryonic thermal index (ETI) on YOY density in Lakes Annecy (black circles) and Bourget (white circles), according to the G hierarchical GAM (Pedersen et al., 2019).

## 4 DISCUSSION

### 4.1 Dominant role of embryonic thermal conditions

Although the dataset covers a relatively short period of approximately 10 years, our results clearly demonstrate the importance of temperature-based predictors, particularly the embryonic thermal index (ETI), in explaining year-to-year variations in the density of YOY perch. These results support our hypothesis that fine-scale thermal indices based on phenology, such as the ETI, provide a more accurate explanation of recruitment variability than conventional seasonal temperature measurements do. Consistent with this finding, a strong correlation was observed between the ETI and the density of YOY perch from one year to the next at the end of summer in both lakes. These findings suggest that recruitment success is largely determined by abiotic conditions. Indeed, ETI was the only factor consistently associated with recruitment patterns across both systems. High ETI values, reflecting steady warming with limited day-to-day variability, were associated with higher YOY densities, whereas low ETI values, driven by stagnant or fluctuating thermal conditions, coincided with reduced recruitment. ETI and TEV capture different aspects of thermal variability. TEV measures the magnitude of temperature fluctuations; however, it ignores their direction. In contrast, ETI considers whether temperatures are increasing or decreasing, thereby reflecting the thermal trajectory during incubation. This distinction likely explains why, unlike TEV, ETI explained recruitment variability, as embryonic development depends more on consistent warming than on variability alone.

In Lake Bourget, ETI explained 44% of the variance in YOY density, and in Lake Annecy, it explained up to 88%, highlighting the dominant role of embryonic thermal conditions in shaping recruitment dynamics. These findings are consistent with those of earlier studies on yellow perch showing that egg mortality increases under stagnant or decreasing temperatures during embryonic development (Clady, 1976; Guma’A, 1978). Incubation near optimal temperatures has also been linked to larger larval size at hatching and improved early survival (Wang, 1994). More broadly, spring temperatures have been shown to positively influence both larval and juvenile densities (Dembkowski et al., 2017; Pope et al., 1996; Ward et al., 2004), in line with our results for Eurasian perch. Hierarchical modelling further confirmed that this positive thermal effect is shared across lakes and outweighs system-specific differences. The most parsimonious models consistently supported a common thermal recruitment relationship, indicating that warming during the incubation phase benefits perch recruitment irrespective of local environmental conditions. Furthermore, the difference in explained variance by ETI between Lake Bourget (44%) and Lake Annecy (88%) could be due to the lower percentage of YOY perch in the upper layer of Lake Bourget (74%) compared to Lake Annecy (93%). Consequently, the proxy for YOY perch density would be less reliable at Lake Bourget, resulting in a lower response to the ETI.

During the spawning period, thermal stratification is still weak, allowing wind-driven mixing or localized upwelling in littoral areas (Gillet & Dubois, 1995), which can bring colder water to the surface and increase embryonic mortality (Clady, 1976; Newsome & Aalto, 1987). This likely explains the abrupt temperature drops observed in some years. Wind-driven currents may also displace egg ribbons into suboptimal habitats (Čech et al., 2011; Clady & Hutchinson, 1975), further increasing mortality. Although we did not distinguish between mechanical and thermal effects, it is likely that both contribute to variability in interannual recruitment. Conversely, rapid spring warming can increase recruitment by accelerating egg development and reducing exposure time to predation, dispersal, and physical disturbance (Clady, 1976; Clady & Hutchinson, 1975; Newsome & Aalto, 1987). It may also limit thermal stress, reduce developmental abnormalities (Hokanson & Kleiner, 1974), and improve match–mismatch dynamics with zooplankton peaks (Busch et al., 1975; Pitlo, 2002).

### 4.2 Timing of spring warming

Some studies have reported that perch recruitment is strongest when warm conditions persist throughout both the embryonic and larval stages (Clady, 1976; Kipling, 1976). According to the growth hypothesis (Anderson, 1988), extended warm conditions from spring to summer can influence year–class strength, as observed in *P. fluviatilis* and *P. flavescens* populations (Craig & Kipling, 1983; Dembkowski et al., 2017). Geographically, Lakes Annecy and Bourget are in the southern part of the species’ natural distribution, and summer surface temperatures therefore rarely decrease below 14 °C, consistently exceeding the lower growth threshold of 13.5 °C (Power & Van Den Heuvel, 1999). Spring–summer temperatures were generally above levels limiting larval and juvenile development. These observations indicate that posthatching thermal conditions were unlikely to constrain recruitment.

Taken together, these results show that recruitment depends on not only temperature magnitude but also the timing and stability of warming. In Lake Bourget, a recent shift towards earlier and more rapid spring warming was observed, with spawning occurring 14–20 days earlier than average in March 2020 and 2022. In both years, temperatures rapidly reached optimal values (∼11 °C) before abruptly declining, leading to low ETI values and reduced YOY densities. These results suggest that early but unstable warming events can negatively affect recruitment, likely by exposing embryos to renewed cold stress (Van De Hey et al., 2013) or disrupting developmental synchrony (Cushing, 1990; Durant et al., 2007). Thus, beyond overall warming, the temporal dynamics of spring temperature appear critical for recruitment success, and increasing climate variability may fundamentally alter recruitment patterns.

### 4.3 Thermal conditions during the larval stage

During the larval phase, a significant effect was detected only in Lake Annecy. Larval thermal amplitude (LTA), which represents the temperature range experienced during this period, was the only index significantly related to YOY density, with higher amplitude associated with lower recruitment. A similar negative pattern was also observed for Annecy for the larval thermal index (LTI), although this relationship was marginally nonsignificant.

LTA was also significantly correlated with ETI, indicating that years with low embryonic thermal conditions tend to be followed by higher thermal variability during the larval stage. In practice, low ETI values were consistently associated with both reduced YOY density and elevated LTA, as surface temperatures eventually reached similar levels (∼20 °C) by June in all years. Notably, high LTA values often coincided with years including modelled temperature data, which may slightly inflate thermal variability due to smoothing effects. Conversely, low LTA values corresponded to years with already warm and stable conditions at the beginning of the larval phase, which appeared more favourable for recruitment.

### 4.4 Role of trophic factors and food availability

Temperature also influences perch energetics and feeding dynamics by regulating both food intake and prey phenology, potentially leading to match–mismatch dynamics between larvae and zooplankton (Craig, 1978, 1987). However, the critical window for prey availability is short (Wang & Eckmann, 1994), and our dataset lacked high-frequency zooplankton data, preventing a precise evaluation of this mechanism. Indeed, zooplankton abundance was not a significant predictor of YOY density in either lake in our analysis. However, laboratory work has demonstrated that prey size, type, and abundance strongly affect larval and juvenile perch growth and survival (Arts & Sprules, 1989; Graeb et al., 2004; Romare, 2000). Slow growth during early summer can extend the period of vulnerability to predation, decreasing survival (Bailey & Houde, 1989; Miller et al., 1988). For instance, in Lake Annecy, zooplankton do not appear to limit growth, possibly because of sufficient resource availability in this oligotrophic system (Perga et al., 2009; Perrier et al., 2012). In contrast, other systems have shown strong links between prey availability and perch dynamics (Dubois et al., 2008). In our study, the zooplankton indices showed no temporal trend but were correlated with nutrient concentrations, suggesting that they may act as proxies of trophic status rather than direct drivers of recruitment (Alric et al., 2013).

### 4.5 Influence of adult stock and predation

Adult stock effects were difficult to interpret. In Lake Bourget, the Gen index (spawning stock for year n+1) was negatively associated with YOY density, whereas the Pred index (potential cannibalism) showed a nonsignificant positive trend; these patterns likely reflect methodological limitations rather than true biology. Estimates of adult abundance were derived from gillnet surveys, which are prone to selectivity bias and undercatch (Olin et al., 2009). The link between fishing mortality and commercial fishing was also not considered. Neither index was significant in Lake Annecy.

Predation affects perch recruitment, including cannibalism (Craig, 1978, 2000; Dembkowski et al., 2017; Gillet & Dubois, 2009). However, in our systems, the effects of adult stock and cannibalism appear to be secondary, partly due to data limitations. Other predators (e.g., pike) were excluded due to lack of detailed data; although predation effects can vary across systems, they can sometimes indirectly promote recruitment by reducing competition (Dembkowski et al., 2017). Future work should aim to improve adult stock indicators to clarify the roles of cannibalism and spawning stock on YOY dynamics.

### 4.6 Methodological limitations and contributions to the field

We were unable to detect any correlations between, on the one hand, the availability of zooplankton and parameters related to adult stocks and, on the other hand, the density of YOY perch. Furthermore, differences in data quality and resolution are reflected in contrast with temperature-based indices. Whereas thermal variables were derived from continuous, high-frequency measurements, biological variables were based on more limited and indirect estimates. The zooplankton data were collected at a much lower temporal resolution (monthly), potentially causing critical short-term dynamics to be overlooked. Adult perch abundance was inferred from gillnet surveys, which are subject to selectivity bias and undercatch. These limitations likely reduced the ability of the biological variables to explain the variability in recruitment.

This study also demonstrates the value of long-term hydroacoustic monitoring. Such surveys are rare in freshwater systems, especially across multiple lakes over decades (Pollom & Rose, 2016). Our work shows that these techniques provide robust ecological insights and a unique perspective on the environmental drivers of fish recruitment in lakes.

### 4.7 Conclusions and perspectives

This study revealed that the success of perch recruitment depends primarily on the temperature conditions experienced during the embryonic stage. Warming without significant day-to-day temperature fluctuations during this critical period results in higher densities of YOY perch. By showing that fine-scale temperature indicators based on phenology perform better than conventional seasonal averages do, our results emphasize the importance of aligning environmental descriptors with biological calendars. In the context of current global climate change, this mechanistic understanding provides a valuable basis for anticipating how changes in spring temperature regimes might affect recruitment dynamics in lake ecosystems.

## AUTHOR CONTRIBUTIONS

V.C. led the study, performed the analyses, and drafted the manuscript. C.Gou. co-supervised the project, supported the study throughout and contributed to manuscript preparation and writing. J.G. co-supervised the project, secured funding, and contributed to hydroacoustic analyses and writing. C.B. contributed to statistical analyses and manuscript writing. O.A. supported later stages of writing and provided critical revisions. N.S. conducted lake temperature simulations. C.Gil. contributed legacy expertise and background knowledge on Eurasian perch in Lake Geneva. All authors reviewed, edited, and approved the final version of the manuscript.

## ACKNOWLEDGEMENTS

We gratefully acknowledge the UMR CARRTEL field teams, M. Colon, P. Chifflet, J.C. Hustache, and C. Rautureau for acquiring the hydroacoustic data during nightly surveys conducted over a decade under challenging conditions. Hydroacoustic data analysis was performed by M. Colon and C. Rautureau, and we extend our thanks for their valuable help. We also thank the CISALB (Intercommunal Committee for the SAnitation of Lake Bourget), the SILA (Lake Annecy Mixed Syndicate) and the Grand Lac community for providing temperature data as well as the Observatory on LAkes for the data set (http://www6.inra.fr/soere-ola © OLA-IS, AnaEE-France, INRAE, Thonon-les-Bains, Syndicat Mixte du Lac d’Annecy, Comité Intercommunautaire pour l’Assainissement du Lac du Bourget, developed by Eco-Informatics ORE INRAE Team). This work was supported by Pôle ECLA (OFB - INRAE - USMB) with additional support from AnaEE France and OLA (boat and technical facilities).

---

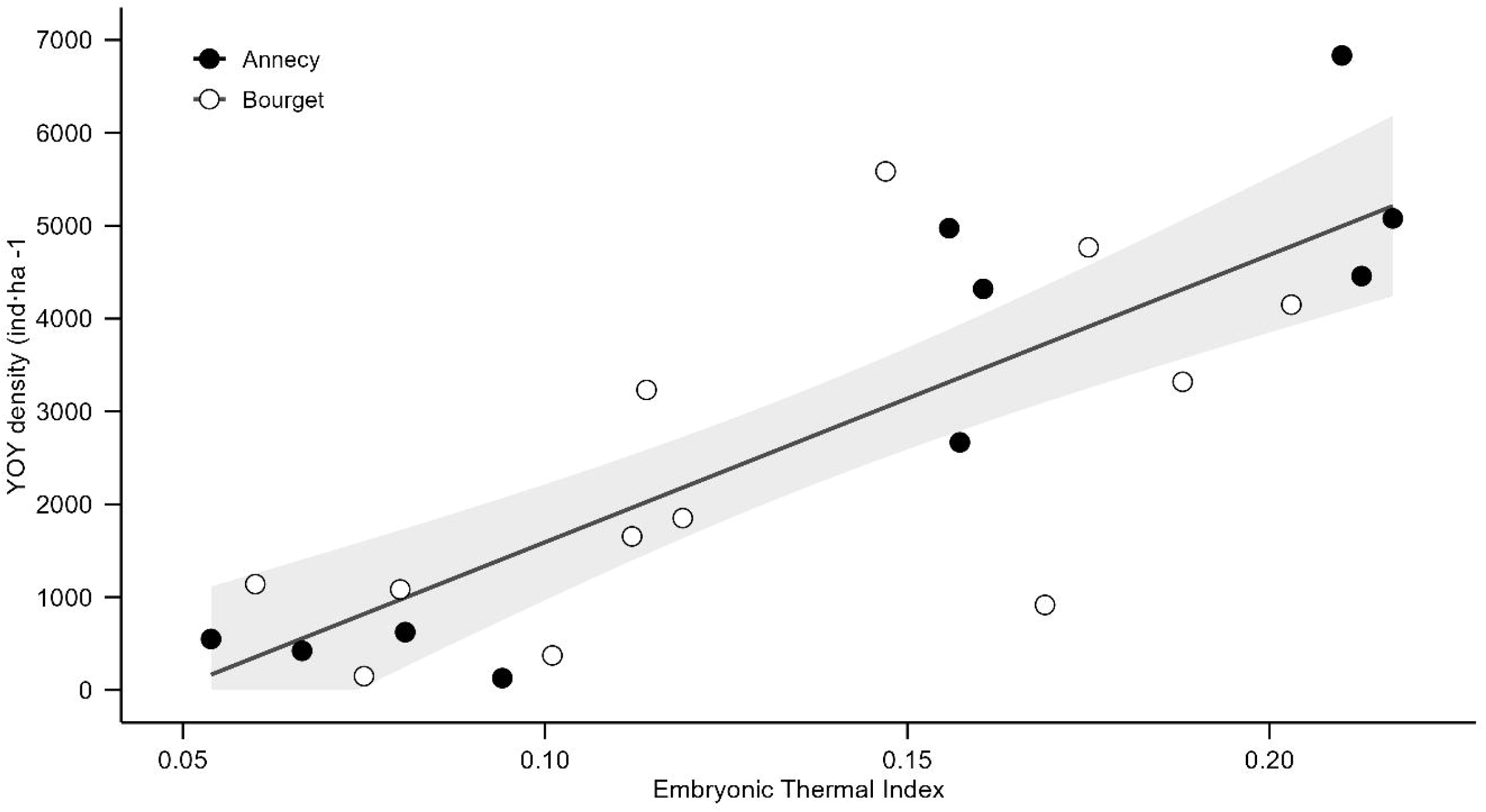

