## Supporting Information for "Early-life temperature drives recruitment success in Eurasian perch (*Perca fluviatilis*) populations"

*S1 - Lake trophic status*

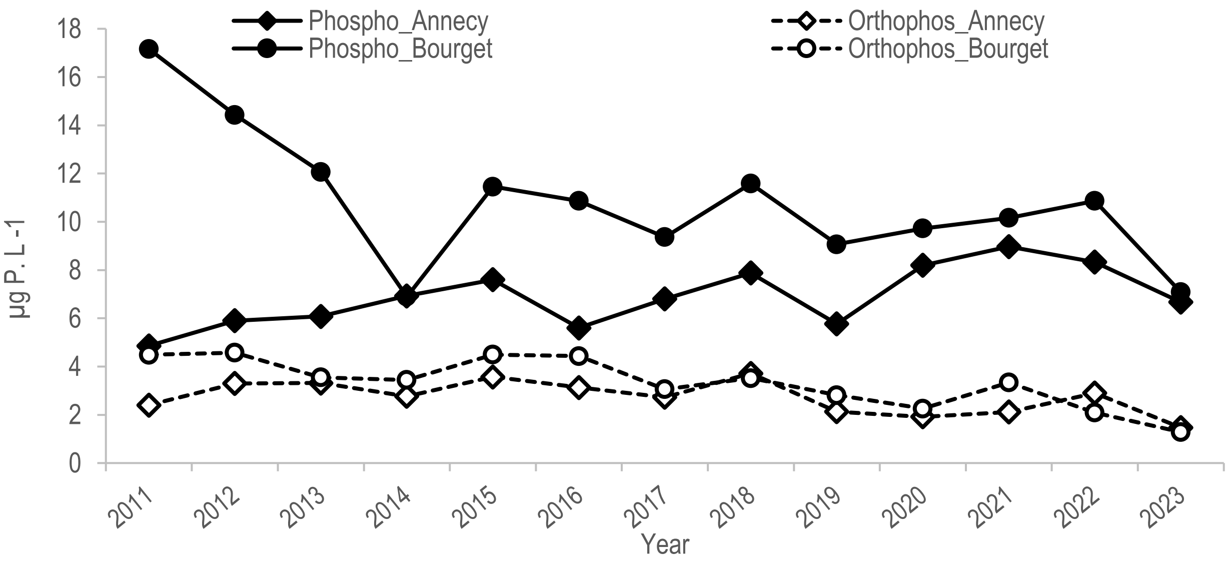


Supplementary Figure 1: Trends for Lake Annecy (diamond) and Lake Bourget (circle) in total phosphorus (full) and orthophosphorus (empty) from April to June between 0 and 15 m depth for the period 2011-2023 (OLA survey data http://www6.inra.fr/soere-ola © OLA-IS, AnaEE-France, INRAE, Thonon-les-Bains, SILA, CISALB, developed by Eco-Informatics ORE INRAE Team). To assess the potential influence of lake trophic status and its long-term evolution on perch recruitment, we selected the mean concentrations of total phosphorus and orthophosphorus between April to June as a proxy. Phosphorus is widely recognized as the most limiting nutrient for primary producers in lake ecosystems and thus plays a key role in structuring trophic dynamics. As such, it can indirectly influence zooplankton abundance at the time of perch larval emergence. During this critical period of first feeding, prey availability is essential for survival. This phosphorus-based index is therefore of dual interest: it reflects both the trophic status of the two lakes and provides an indirect measure of potential prey availability for larval perch.

*S2 - Phenology date determination*

Supplementary Table 1: Theorical date of development phases with E for embryonic phase and L for larval phase. Last two columns, duration, are in days; the second table represents average by lake.

| Year | Lake | Beginning_E | Ending_E | Beginning_L | Ending_L | E_Duration | L_Duration |
|---|---|---|---|---|---|---|---|
| 2012 | Annecy | 27-april | 22-may | 12-may | 25-june | 25 | 44 |
| 2013 | Annecy | 30-april | 12-june | 16-may | 15-july | 43 | 60 |
| 2014 | Annecy | 07-april | 10-may | 23-april | 10-june | 33 | 48 |
| 2016 | Annecy | 28-april | 20-may | 13-may | 21-june | 22 | 39 |
| 2018 | Annecy | 18-april | 14-may | 01-may | 15-june | 26 | 45 |
| 2019 | Annecy | 18-april | 20-may | 03-may | 21-june | 32 | 49 |
| 2020 | Annecy | 06-april | 29-april | 20-april | 31-may | 23 | 41 |
| 2021 | Annecy | 21-april | 24-may | 08-may | 25-june | 33 | 48 |
| 2022 | Annecy | 14-april | 18-may | 29-april | 19-june | 34 | 51 |
| 2023 | Annecy | 22-april | 16-may | 07-may | 17-june | 24 | 41 |
| 2011 | Bourget | 01-april | 22-april | 17-april | 24-may | 21 | 37 |
| 2012 | Bourget | 28-april | 18-may | 12-may | 19-june | 20 | 38 |
| 2013 | Bourget | 14-april | 09-may | 30-april | 10-june | 25 | 41 |
| 2014 | Bourget | 31-march | 05-may | 15-april | 06-june | 35 | 52 |
| 2015 | Bourget | 12-april | 08-may | 27-april | 09-june | 26 | 43 |
| 2016 | Bourget | 11-april | 17-may | 27-april | 18-june | 36 | 52 |
| 2018 | Bourget | 15-april | 14-may | 28-april | 15-june | 29 | 48 |
| 2019 | Bourget | 15-april | 22-may | 30-april | 23-june | 37 | 54 |
| 2020 | Bourget | 19-march | 16-april | 06-april | 18-may | 28 | 42 |
| 2021 | Bourget | 18-april | 14-may | 02-may | 15-june | 26 | 44 |
| 2022 | Bourget | 25-march | 25-april | 12-april | 27-may | 31 | 45 |
| 2023 | Bourget | 09-april | 10-may | 25-april | 11-june | 31 | 47 |

| Lake | Mean Beg_E | Mean End_E | Mean Beg_L | Mean End_L | Mean E_dur | Mean L_dur |
|---|---|---|---|---|---|---|
| **Annecy** | 19-april | 18-may | 04-may | 19-june | 29,5 ± 6,6 | 46,6 ± 6,1 |
| **Bourget** | 08-april | 07-may | 24-april | 08-june | 28,75 ± 5,5 | 45,25 ± 5,5 |

*S3 - Perch spawning phenology in Lake Geneva*

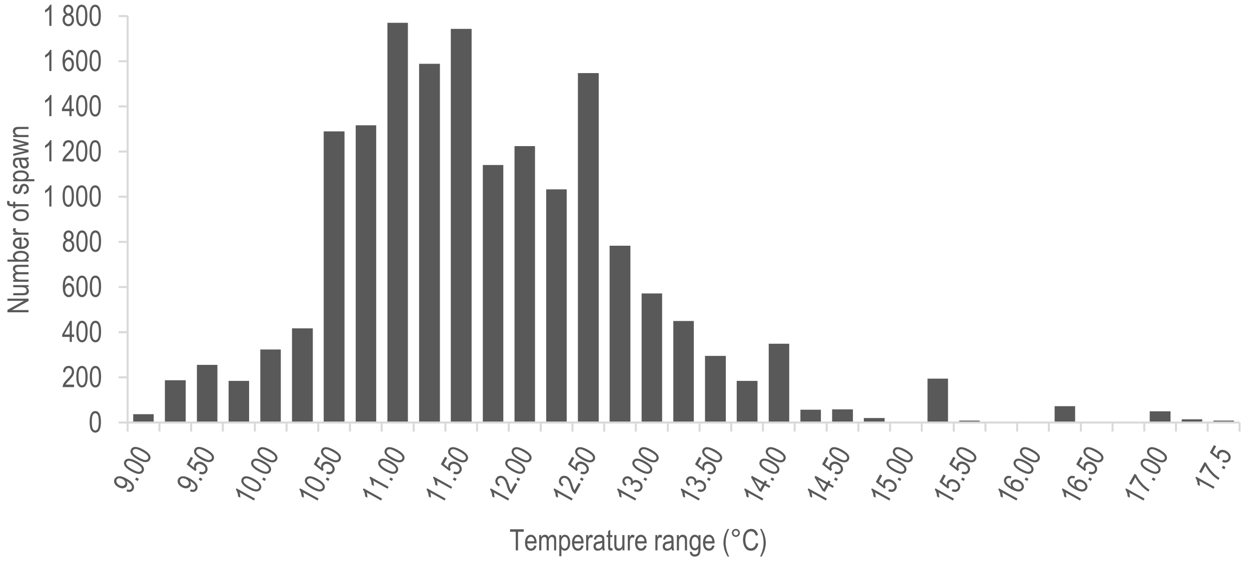


Supplementary Figure 2: Perch phenology in Lake Geneva. Distribution of the 17.181 spawns collected on artificial spawning ground from 1986 to 2022 across temperature range (Gillet & Dubois, 2007; Goulon et al., 2024).

*S4 - Temperature data*

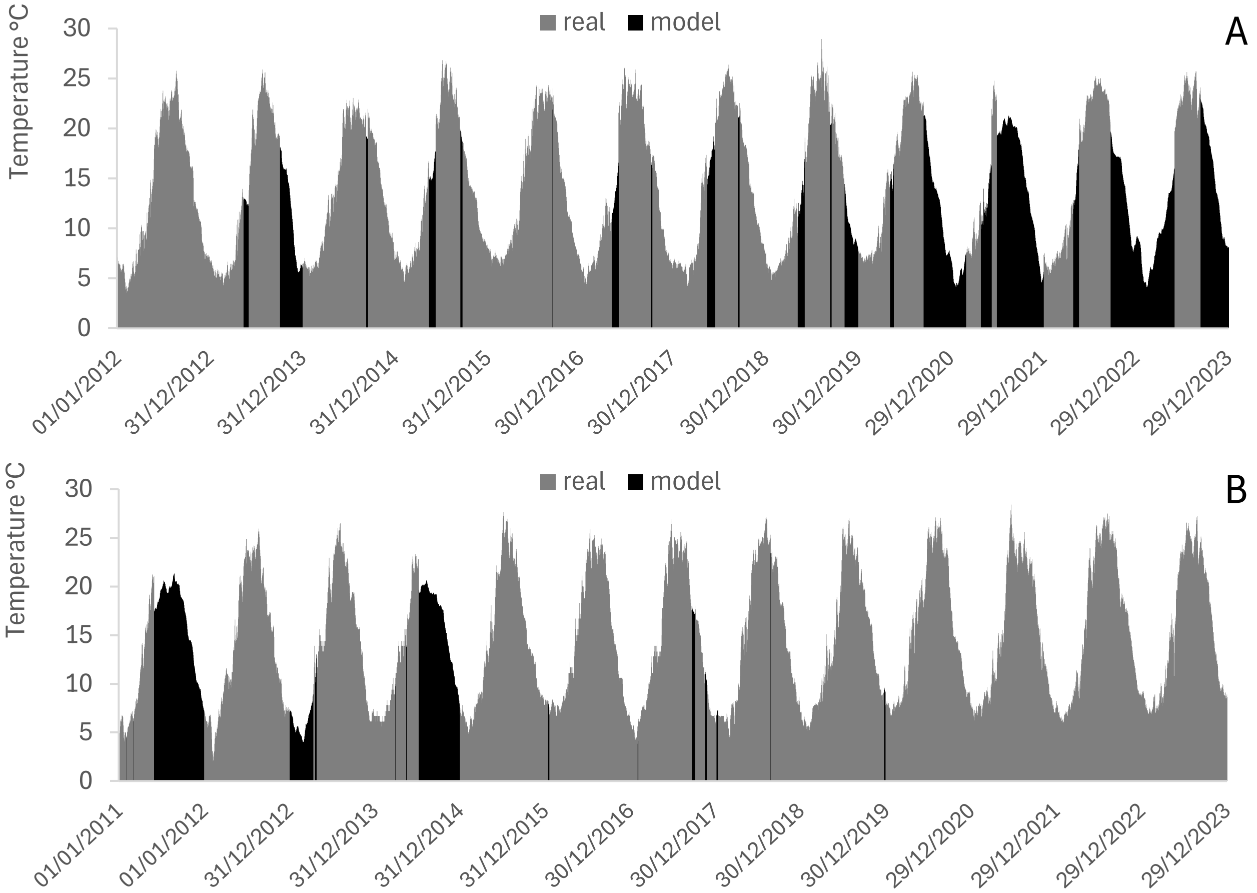


Supplementary Figure 3: Type of temperature data used, with in grey, the real data measured on site, and in black, the modelled data, for Lake Annecy (A) and Lake Bourget (B).

*S5 - Example of ETI calculation (Lake Annecy, 2012)*

To illustrate the calculation of the Embryonic Thermal Index (ETI), we provide an example based on the 2012 incubation period in Lake Annecy (Supplementary Table 2).

Supplementary Table 2 :  Incubation period temperatures, Annecy 2012

| *Day of incubation* | ° C | TV |
|---|---|---|
| *1* | 10,49 | |
| *2* | 11,58 | 1,085 |
| *3* | 10,53 | -1,049 |
| *4* | 10,21 | -0,320 |
| *5* | 10,60 | 0,392 |
| *6* | 11,04 | 0,440 |
| *7* | 11,64 | 0,601 |
| *8* | 12,27 | 0,631 |
| *9* | 12,02 | -0,255 |
| *10* | 12,34 | 0,319 |
| *11* | 13,06 | 0,722 |
| *12* | 12,44 | -0,619 |
| *13* | 13,03 | 0,587 |
| *14* | 13,92 | 0,890 |
| *15* | 14,86 | 0,944 |
| *16* | 14,60 | -0,264 |
| *17* | 11,09 | -3,506 |
| *18* | 12,64 | 1,547 |
| *19* | 13,43 | 0,792 |
| *20* | 13,04 | -0,388 |
| *21* | 13,63 | 0,591 |
| *22* | 13,91 | 0,282 |
| *23* | 14,25 | 0,337 |
| *24* | 14,96 | 0,712 |
| *25* | 14,96 | -0,006 |
| *26* | 14,42 | -0,533 |

ETI is computed as the sum of daily temperature variations (TV), defined as the difference between consecutive daily mean temperatures:

$$ETI = \sum_{i=1}^{n} TV$$

During this incubation period (27 April to day 26), water temperature increased overall from 10.49 °C to 14.96 °C (Supp. Figure 4). However, this warming trajectory included several short-term fluctuations, including a marked cold event on day 17 (TV = − 3.5 °C).

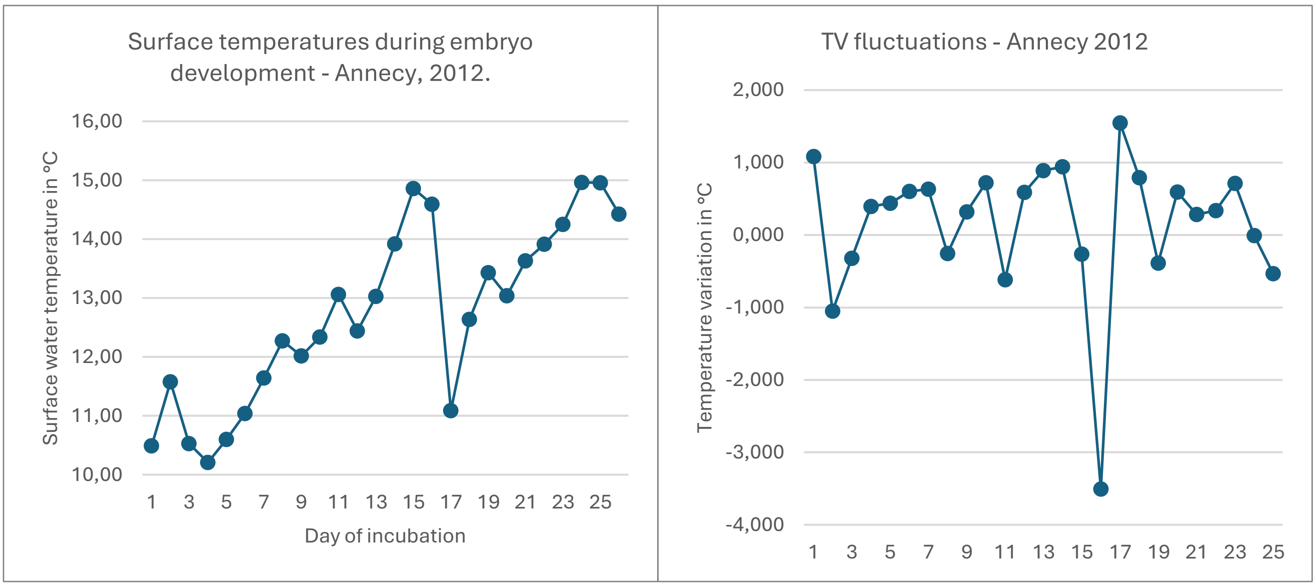


Supplementary Figure 4: Temperature profile during incubation and TV fluctuations in Annecy 2012.

This example highlights that, although ETI is based on the sum of daily temperature changes, it does not simply reflect the net difference between initial and final temperatures. Instead, it integrates the sequence of warming and cooling events throughout the incubation period. In particular, cooling events reduce the cumulative index, thereby capturing unfavourable thermal conditions that would not be reflected by a simple temperature difference.

*S6 - Validation of ETI calculation method*

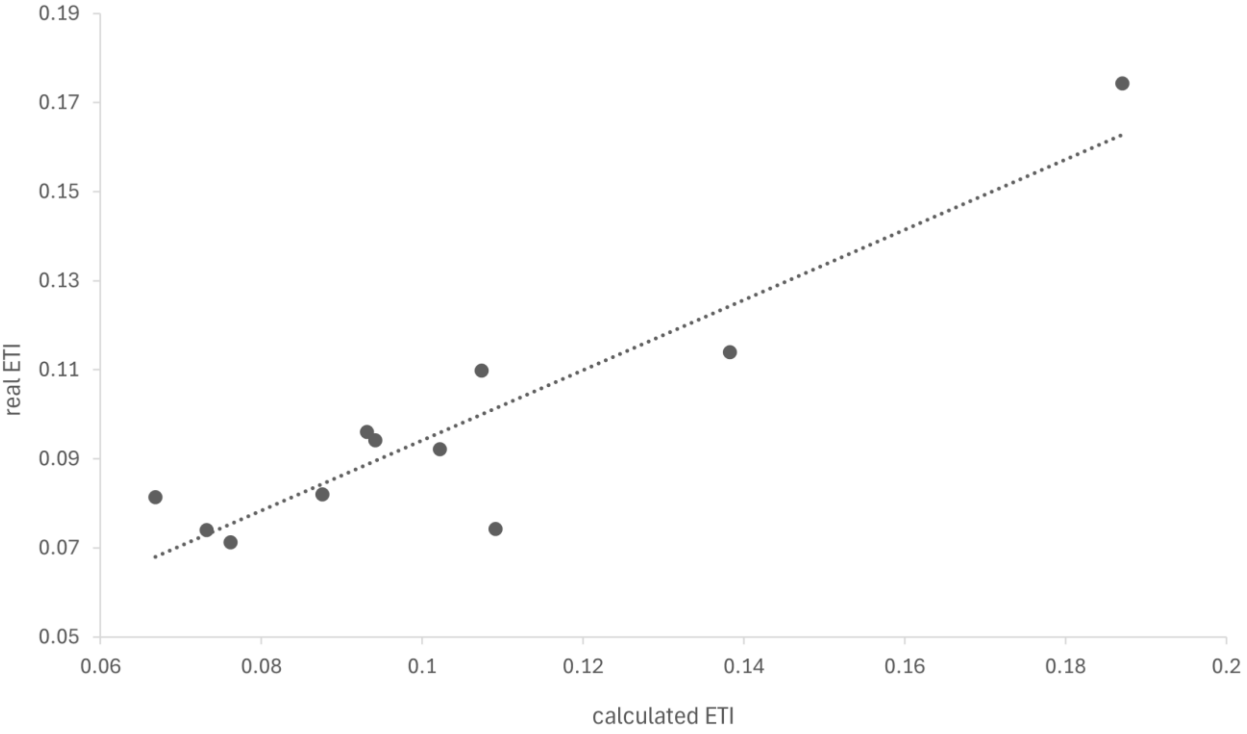


Supplementary Figure 5: Comparison of ETI values derived from two calculation methods: one based on observed phenology and the other on a temperature-based prediction method using thresholds of 10.5°C and 13.3°C. To validate index calculation methods and the theoretical reproduction/development period, we compared the ETI values applied to Lake Geneva. On this lake, we have empirical spawning data and modelled surface temperatures (Sharaf et al., 2023). It was therefore possible to calculate two different sets of indices. The first set of ETIs was calculated by taking measured values as the start and end of spawning, recorded during monitoring from 2011 to 2022 to correspond to our study period on lakes Annecy and Le Bourget. The second set of ETIs was calculated as explained in this article, i.e. using the 10.5°C and 13.3°C thresholds as the start and end of spawning. No significant difference was found between the two approaches (*p*-value = 0.577). The linear model relating observed and predicted ETI values was highly significant (*p*-value = 6.81e-05 ***), with an adjusted $R^2$ of 0.82. A two-factor Anova (calculation method * year) was also performed. Only the factor year shows significant differences between ETI values (*p*-value = 0.006**). No significant differences were observed between ETI values calculated on real phenology data and on spawning periods determined based on the surface temperature of Lake Geneva. The ETI, being the most complex index, was the only one validated against observed spawning data. Its consistency supports the assumption that simpler indices are also reliable.

*S7 - Wind parameter during the embryonic and larval phases*

For the embryonic phase, wind is another parameter in addition to temperature that can explain the success or failure of incubation (Bourinet et al., 2023). Strong wind speeds and current can destroy or displace eggs to a less hospitable habitat (Clady, 1976). We used daily wind speed data available on the period 2011-2023 from a station at Meythet (74 - Haute-Savoie, France) which is relatively close to Lake Annecy (4 kilometres) but situated at 30 kilometres from Lake Bourget. However, even though the two valleys are oriented in the same direction, the winds conditions differ between the lakes, making the wind proxy for Lake Bourget is therefore less than optimal. An Embryonic Wind Condition index, EWC, has nevertheless been constructed for each lake. It corresponds to the mean value of wind speed in metres per second during the embryonic phase. Because the impact of strong winds and flows on larvae can be significant, with displacement into an early pelagic environment, depth displacement, destruction, etc., a Larval Wind Condition index, LWC, has also been constructed for each lake. It corresponds to the mean value of wind speed in metres per second during the larval phase.

*S8 - Zooplankton abundance during the larval phase*

Zooplankton abundance indexes were build for the main taxa predated during the larval stage : Rotifers, Copepods and Cladocera (Craig, 2000; Masson et al., 2001; Wang, 1994). These indexes use data from the Observatory of LAke (OLA) monitoring surveys (http://www6.inra.fr/soere-ola © OLA-IS, AnaEE-France, INRAE, Thonon-les-Bains, SILA, CISALB, developed by Eco-Informatics ORE INRAE Team ; Rimet et al., 2020). Zooplankton are sampled along a vertical tow beginning at 50 m from the surface, using a 200-µm plankton net for crustaceans and a 64-µm plankton net for rotifers. Samples were fixed on the boat using 5% buffered formaldehyde. Identification and counting of crustaceans were performed in a subsample of a known volume of the net sample using a light microscope at x10 magnification. The results were given in number of individuals per square metre (Rimet et al., 2020). Zooplankton abundance indexes were the average density of each prey category of perch larvae calculated over the period of the larval stage which gives an average of the available target zooplankton per square meter of water column (0-50m). Zooplankton Abundance Index - Rotifers (zai_rot) represents the average abundance of rotifers. Zooplankton Abundance Index - Copepod (zai_cop) represents the abundance of immature copepods (nauplii and copepodites) and adult copepods during larval phase. Zooplankton Abundance Index - Cladoceran (zai_clad) represents the abundance of cladoceran taxa and finally, Zooplankton Abundance Index - All (zai_all) is the sum of the three previous indexes, every prey abundance during larval phase.

*S9 - Spawner stock and cannibalism*

The monitoring of fish populations in peri-alpine lakes has enabled the calculation of the biomass of adult perch caught per 1,000 m² of net set every year during scientific surveys (Rimet et al., 2020). The biomass value obtained for year n at the end of the summer season is used to test the effect of cannibalism on the density of YOY perch in year n. The same value is also used to test the effect of the spawning stock for year n + 1 (Pred index), since the adults of year n will be the spawners for the following year (Gen index).

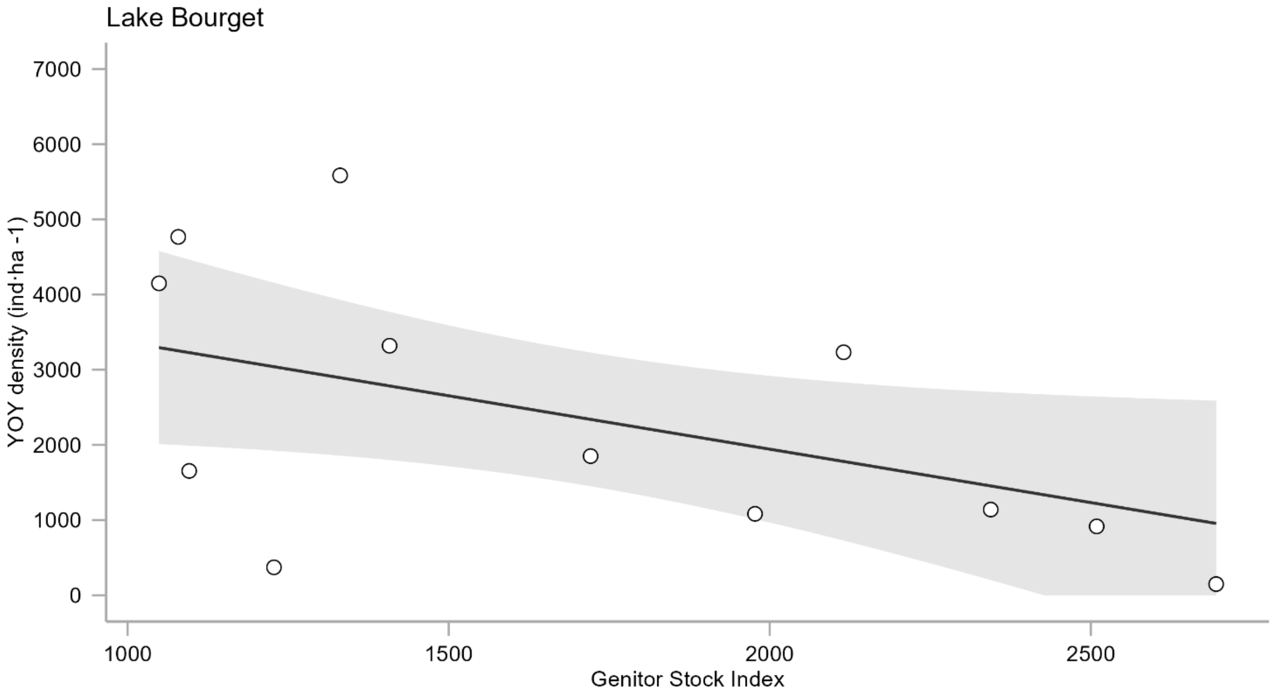


Supplementary Figure 6: Effect of the stock of Genitor index (Gen) on YOY density in Lake Bourget, single covariate GAMs.
